# SEEK-VEC: Augmenting topic modeling with spectral ensemble learning

**DOI:** 10.64898/2025.12.12.693799

**Authors:** Rebecca Danning, Zheng Tracy Ke, Rong Ma, Xihong Lin

**Affiliations:** Center for Genomic Medicine, Massachusetts General Hospital; Stanley Center, Broad Institute of MIT and Harvard; Department of Statistics, Harvard University; Department of Biostatistics, Harvard T.H. Chan School of Public Health; Eric and Wendy Schmidt Center, Broad Institute of MIT and Harvard; Department of Data Science, Dana-Farber Cancer Institute

## Abstract

Count data are ubiquitous across many applications in which understanding latent patterns is of interest. Topic modeling is a powerful tool for detecting latent structure in count data. However, standard topic modeling methods are often constrained by their restrictive assumptions, susceptible to noise, and sensitive to misspecification of the number of topics. Here, we introduce SEEK-VEC (Spectral Ensembling of topic models with Eigenscore for K-agnostic Vocabulary Embedding and Classification), an ensemble topic modeling framework that integrates insights from multiple candidate topic models through a spectral ensembling procedure. SEEK-VEC produces a meta-structure matrix containing prioritization scores and grouping scores that enable variable classification, interactive pattern discovery, and model diagnostics. Through simulations, we demonstrate that SEEK-VEC augments the performance of standard topic models for identifying important vocabulary words and understanding the relationships among them, particularly when signal strength is weak. We apply SEEK-VEC to the MADStat dataset of statistical abstracts and demonstrate its utility for evaluating the proposed interpretation of a topic model.

## 1 Introduction

Topic modeling is a statistical approach in text analysis that aims to identify a small set of latent topics underlying a large collection of documents.^1^ Topic modeling is typically used on integer data representing counts of words per document, but can also be applied to non-text integer data settings. For example, topic modeling has been used in on a wide variety of biomedical data, such as microarrays,^2^ genome annotation,^3^ and MRI data.^4^ The “topics” found from biomedical data represent latent biological concepts that can yield insight into underlying structure.^5^ Additionally, the output of a topic model can be used as a feature selection tool separating signal-relevant words from uninformative words, even in non-text settings without a conventional “vocabulary.”^6^

However, there are several necessary considerations when using topic modeling for analyzing data. First, most existing topic modeling methods require specifying the number of topics *K* in advance for implementation.^7–10^ But consistently estimating *K* from the data remains a challenging task in real-world applications.^7^ Several data-driven approaches have been developed for estimating *K*, include analyzing the singular values of the data,^11^ perplexity-based cross-validation techniques,^12^ and model-fitting criteria such as BIC,^13^ yet so far there is no consensus on their relative performance and reliability in different applications. Alternatively, *K* can be chosen in part due to the interpretability of the topics based on subject-matter expertise,^1^ but this leaves the model open to subjectivity bias.

Second, many topic modeling methods rely on an “anchor word” assumption, or separability assumption, to ensure model identifiability.^14,15^ For each topic, there should be at least one word – an anchor word – that identifies the topic, meaning that it appears exclusively in one topic and not others.^1,16^ While this condition is usually satisfied in text data,^7^ it is often not the case for non-text applications. For example, statistical analysis of the transcripts from meetings of the Federal Open Market Committee discussing monetary policy strategy rejected the null hypothesis that the corpus contained anchor words.^17^ As such, standard topic modeling algorithms may underperform in non-text applications.

To address these limitations, in this paper we present SEEK-VEC (Spectral Ensembling of topic models with Eigenscore for K-agnostic Vocabulary Embedding and Classification), an ensemble machine learning framework for vocabulary classification, pattern discovery, and model diagnostics that builds on the output from standard topic modeling procedures. SEEK-VEC is particularly valuable in challenging scenarios where the anchor word assumption is violated, and its performance is more stable with respect to hyperparameter selection. It leverages the performance-improving properties of ensemble learning strategies^18,19^ and is a flexible framework for enhancing any existing topic modeling algorithm. Here, we focus on combining our ensemble approach with Topic-SCORE,^1,7^ a state-of-the-art method for topic modeling that uses singular value decomposition to facilitate fast estimation of the low-rank topic matrix with theoretical guarantees. Demonstration of its utility for augmenting alternate topic modeling methods is shown in Supplemental Fig. A1b.

SEEK-VEC produces two complementary sets of scores. Prioritization scores distinguish important variables (or words) from less informative ones, while grouping scores quantify the strength of relationships among variables. Together, these scores enable downstream applications such as variable classification, interactive pattern discovery, and model diagnostics. Through extensive numerical simulations, we show that SEEK-VEC consistently augments existing topic modeling methods across various tasks under realistic noisy data conditions. We then apply SEEK-VEC to a dataset of statistical abstracts to demonstrate its utility for evaluating the output from a proposed topic model. The R package implementation of SEEK-VEC is available at the GitHub repository: https://github.com/rdanning/seekvec.

## 2 Results

### 2.1 Overview of SEEK-VEC

SEEK-VEC builds on the topic model, a statistical framework designed for count data that aims to uncover the latent dimensions, or topics, of the vocabulary. There are three main steps to SEEK-VEC. First, we fit a topic model for each of a range of candidate topic counts *K*. Second, the output of each topic model is converted into a novel sparse representation called the *hallmark structure matrix* that captures the latent embedding information from the model without relying on *K*. Finally, these representations are combined into a consensus-weighted embedding using a spectral ensembling procedure. The diagonal values of this output matrix can be used to prioritize the vocabulary with respect to how much signal it contains of the underlying topic structure, and the off-diagonal values can be used to understand the grouping structure among the vocabulary. SEEK-VEC is summarized in Fig. 1a, and additional details of its implementation are given in the Methods section A.1.

**Figure 1:**
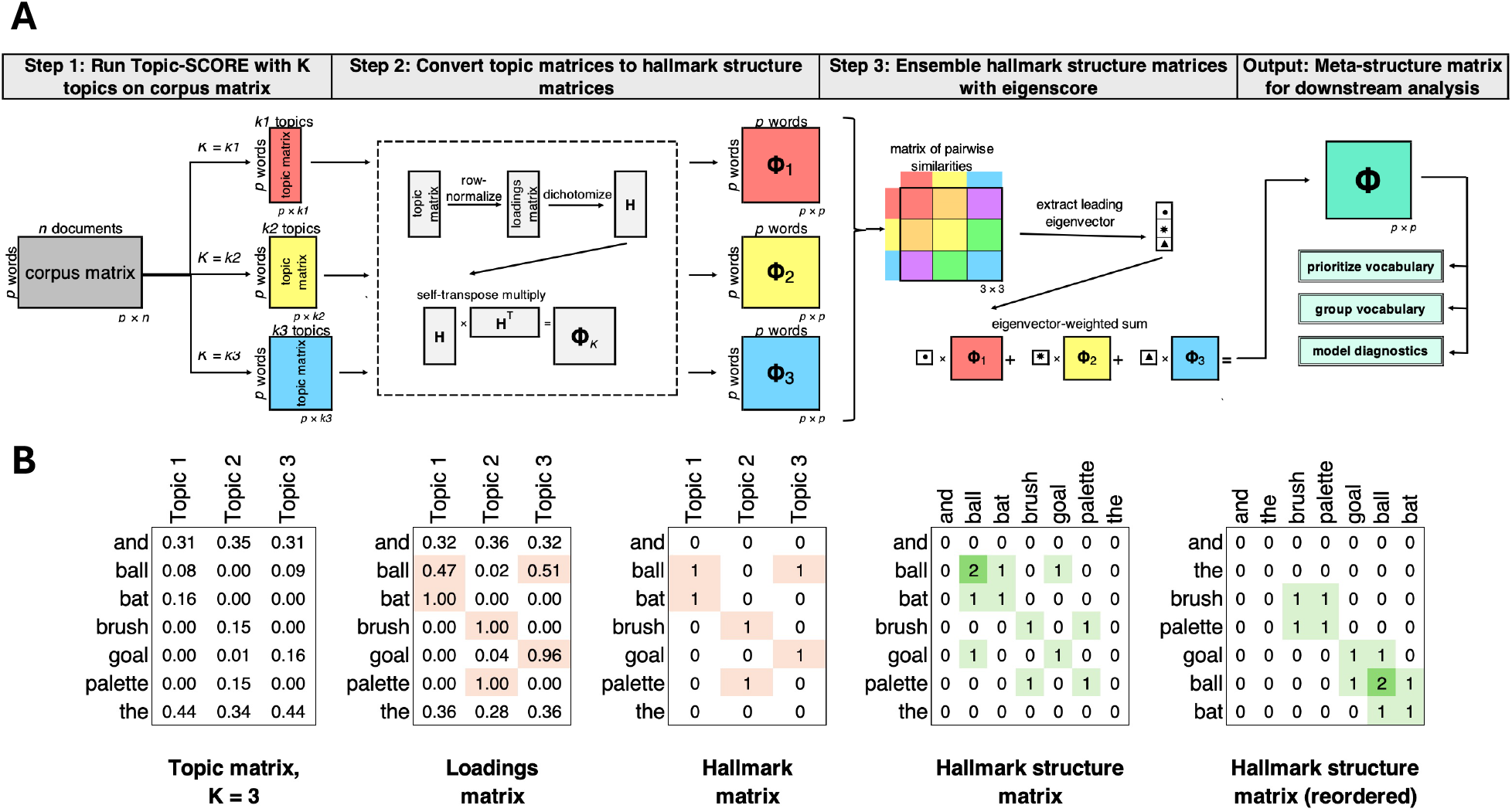
(a) Overview of SEEK-VEC. In the above example, there are three candidate values of *K*: = {*k*1, *k*2, *k*3}. In step 1, topic modeling is run on the input corpus matrix for each candidate value of *K* to yield a collection of ℝ^*p*×*K*^ topic matrices. In step 2, each topic matrix is row-normalized to a topic loadings matrix and then converted into a ℝ^*p*×*p*^ hallmark structure matrix. In step 3, these hallmark structure matrices are ensembled using a spectral ensembling procedure. The output is the meta-structure matrix 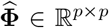. (b) Toy example demonstrating the procedure for converting a topic matrix into a hallmark structure matrix. The topic matrix is the direct output of a topic modeling algorithm, where each column represents a distribution on the vocabulary and sums to 1. The loadings matrix is the row-normalized version of the topic matrix. Notably, words such as “is” and “the”, which are common to all topics, have higher values in the topic matrix but relatively low values in the loadings matrix. The hallmark matrix is a dichotomized version of the loadings matrix. This represents the standard topic modeling procedure of choosing vocabulary words that are representative of each topic. The hallmark structure matrix is the hallmark matrix multiplied by its transpose. Words with a value of 0 along the diagonal (e.g. “is” and “the”) are not representative of any topic. Words with a 1 along the diagonal identify a single topic (e.g. “brush” for Topic 2). Words with a value of 2 along the diagonal link topics (e.g. “ball” links Topic 1 and Topic 3). The values on the off-diagonal indicate the number of topics for which two words are jointly representative. The reordered hallmark structure matrix demonstrates how block structure can be used to recover topics and understand meta-relationships among the topics.

#### Step 1: Run Topic-SCORE on a range of topic counts

Topic modeling algorithms take as input a corpus matrix **D** ∈ ℝ^*p* × *n*^, which contains the counts of each of *p* vocabulary words in each of *n* documents, and a number of topics *K*. Each of the *n* documents is thought to comprise some combination of the *K* topics. The output is a topic matrix **A** ∈ ℝ^*p*×*K*^, where each column represents a probability distribution over the *p* vocabulary words. Each row of **A** then represents a vocabulary word’s position in *K*-dimensional space, with each axis corresponding to a topic. The first step of SEEK-VEC is to run Topic-SCORE^7^ for a collection of candidate values *K*; typically, *K* is a range of consecutive values of *K*. In these simulations, we consider SEEK-VEC with *K* = {3, …, *K*\*}. This step yields a collection of candidate topic matrices **A**_*K*_ for each *K* ∈ *K*. A comparison of alternative topic modeling algorithms can be found in Section A.2.1.

#### Step 2: Convert each topic matrix into a hallmark structure matrix

In practice, interpretation of the topic model is done using the *topic loading matrix*, which is a row-normalized version of the topic matrix, denoted by **B**. The matrix **B** is obtained by dividing each row of **A** by the row sum. This matrix is preferred because it shows the relative weight of each word across the topics, and therefore allows for the interpretation of topics based on the words with the highest loadings.

Once the topic loadings matrix is obtained, the next step in interpretation is choosing representative words for each topic. The representative words for each topic are typically those with loadings above a certain threshold. To replicate this procedure in SEEK-VEC, we next dichotomize the topic loading matrix **B** to a *hallmark matrix* **H** ∈ ℝ^*p*×*k*^, whose entries only take binary values, based on some threshold parameter *t*.

Thinking of **H** as a bipartite graph representing edges between topics and words, the *hallmark structure matrix* **Φ** = **HH** ^⊤^ is equivalent to the one-mode projection network of that graph and summarizes the relationships among the words and topics.^20^ The values along the diagonal of **Φ** indicate the number of topics for which a word is representative. Words with a diagonal value of 1 are only representative for a single topic, and therefore can be used to identify a topic. Words with a diagonal value greater than 1 are meta-informative: these words are representative for more than one topic and therefore indicate a relationship among topics with respect to the vocabulary. Words with a diagonal value of 0 can be disregarded when interpreting the topics. The values on the off-diagonal are also interpretable: the value **Φ**_*ij*_ represents the number of topics for which word *i* and word *j* are both representative, so **Φ**_*ij*_ > 0 indicates a relationship among words with respect to the topics (Fig. 1b). Notably, **Φ** is a *p*×*p* matrix, and thus summarizes information from the *K*-dimensional word embeddings in a form that is free of *K*.

#### Step 3: Ensemble the hallmark structure matrices with eigenscore

Each hallmark structure matrix represents an embedding of the vocabulary in potentially-misspecified *K*-dimensional space. As such, each embedding may contain both signal and noise with respect to the true underlying topic structure. Therefore, we can combine the insights from each model in a way that upweights consensus signal and downweights individual noise using a spectral ensembling method adapted from the *eigenscore*^21^ method (see Section A.1.3). The output of this method is the meta-structure matrix 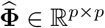 that estimates a consensus embedding of the vocabulary.

The matrix 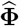 yields key information about the vocabulary and its underlying structure: the diagonal values of the matrix contain *prioritization scores*, which estimate the signal contained in each vocabulary word. These scores can be used to filter the vocabulary: words with zero or near-zero scores likely are uninformative with respect to any topic structure (e.g. “the” in Fig. 1b), whereas words with high scores should be prioritized for downstream analysis. The off-diagonal values of the matrix contain *grouping scores*, where a high grouping score between two words indicates persistent co-occurrence as representative words in the underlying topic models (e.g. “bat” and “ball” in Fig. 1b) and therefore indicate group structure that should be considered in downstream analysis and interpretation.

### 2.2 Advantages of SEEK-VEC and Simulation Results

To test the ability of SEEK-VEC to improve the estimation of the true prioritization and grouping scores, we compare SEEK-VEC, ensembling *K* = 3 to *K* = 12, to each of the component Topic-SCORE matrices. In these scenarios, we simulate a vocabulary of *p* = 2000 words across *K* = 6 topics in *n* = 1000 documents of length *N* = 5000. We simulate 10% of the vocabulary words as identifying words (representative for one randomly-selected topic) and simulate 20% of the words as metainformative words (representative for two randomly-selected topics). In the strong-anchor setting, the representative words have loadings that are 10 × that of the remaining topics. In the weak-anchor setting, the representative words have loadings that are 3 × that of the remaining topics. Results for varying degrees of weakened signal strength are shown in Section A.2.3. In both settings, the remaining 70% uninformative have equal loadings across all topics. Example rows from the simulated loadings matrices are shown in Fig. 2a. The data are simulated under the word frequency heterogeneity setting. In this setting, there is a severe discrepancy in the overall frequency of the vocabulary words, which is common in realistic text settings and is known as Zipf’s Law^22^. We show results from across a range of values of the threshold *t*, where the top *t* proportion of words are selected as representative words; extended results are shown in Section A.2.3.

**Figure 2:**
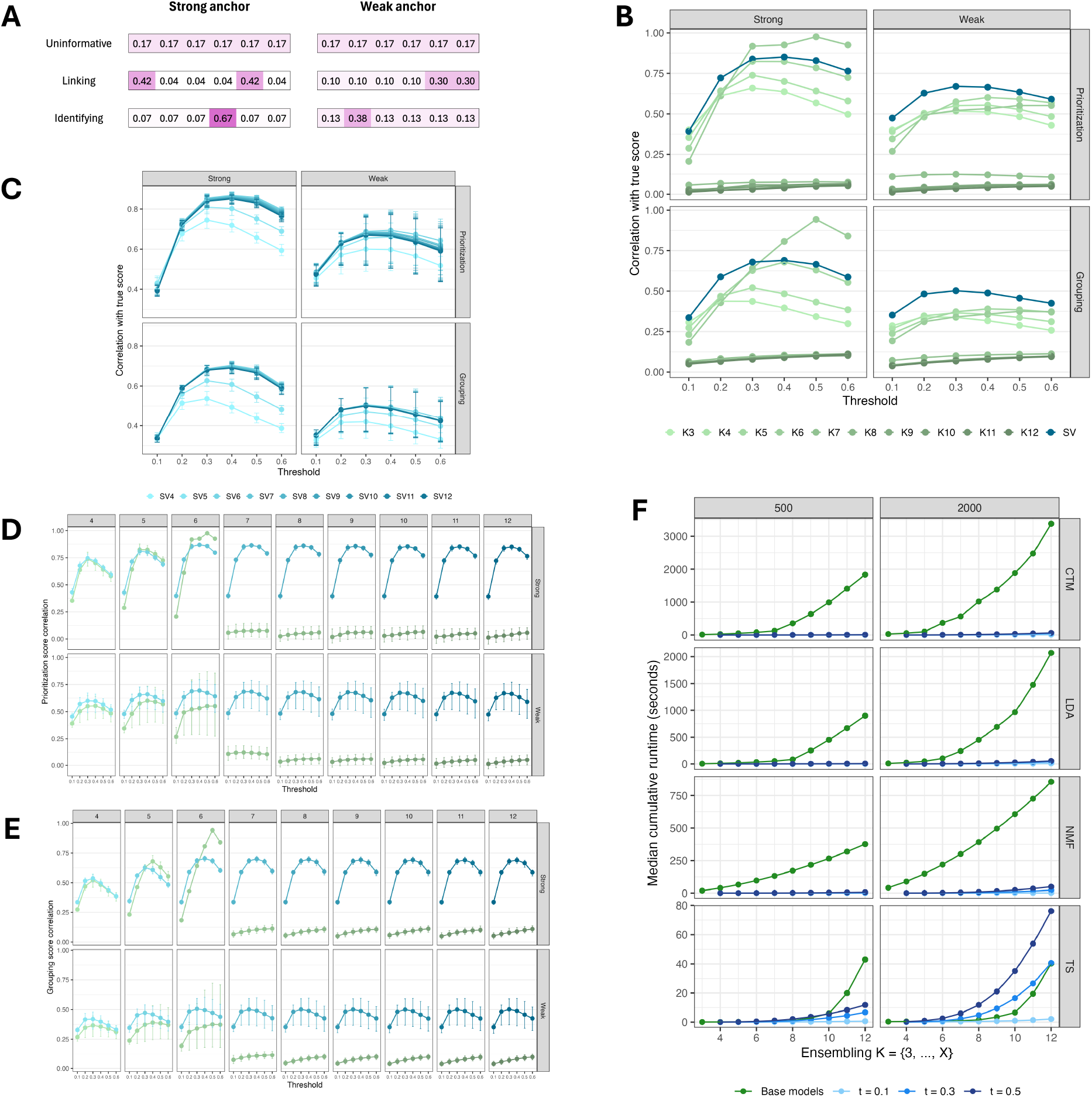
(a) Example rows of from the loadings matrix in the Strong and Weak conditions for each of the three categories of words. (b) Performance of SEEK-VEC compared to the constituent Topic-SCORE models. Prioritization scores estimate how relevant each word is to the underlying topic structure and grouping scores measure how closely-related two words are with respect to the underlying topic structure. The y-axis shows the median across simulations of the estimated scores with the true scores. The x-axis shows the candidate thresholding parameters. (c) Comparison of SEEK-VEC across choice of *K*\*, the upper limit for the ensembling. Error bars represent the 25^*t*^*h* and 75^*t*^*h* percentiles. (d) Direct comparison of the prioritization score correlations for each Topic-SCORE model with its comparative ensemble method; for example, comparing Topic-SCORE with *K* = 7 to SEEK-VEC ensembling *K* = {3, …, 7}. (e) Direct comparison as in (d) for the grouping score correlations. (f) Median cumulative runtimes of the ensembling procedure compared to running the standard topic models for four topic modeling methods. Columns represent different values of *p*. The green lines show the cumulative runtimes of running the base topic model for *K* = {3, …, *X*}, where *X* is the *x*-axis value. The other lines show the additional time taken to ensemble these models with varying values of the threshold *t*.

#### SEEK-VEC more reliably differentiates representative words from uninformative words

The diagonal of the true hallmark structure matrix is equal to the number of topics for which a given word is representative and can be treated as a prioritization score, where words that have higher prioritization scores contain more information about the underlying topic structure. Words that are uninformative, therefore, have a prioritization score of 0, while identifying words have a true prioritization score of 1 and metainformative words have a true prioritization score greater than 1. The top row of Fig. 2b shows the correlations between the true prioritization scores and the and estimated prioritization scores derived from standard Topic-SCORE, as well as SEEK-VEC with *K*\* = 12. In the strong-anchor setting on the left, SEEK-VEC is outperformed by Topic-SCORE with *K* = 6 topics, but improves upon all other Topic-SCORE models – particularly the overspecified (*K* > 6) models – across all threshold values.

Under the weak-anchor condition on the right, the anchor word assumption underlying Topic-SCORE is violated, and thus the performance of all standard models suffers. Notably, though, the performance of SEEK-VEC is better across all threshold levels than that of its component topic models. This indicates that the signal amplification procedure underlying the ensemble process allows for improved estimation even when the assumptions underlying the component models are violated.

#### SEEK-VEC improves estimation of the latent relationships between words

Off-diagonal values in the hallmark structure matrix represent grouping scores between vocabulary words, where two words with a high grouping score are frequently co-identified as hallmark words across candidate topic models. More generally, grouping scores measure the strength of the relationship between two words that persists across candidate models, and patterns of grouping scores can reveal relationships between derived topics. The bottom row of Fig. 2b shows the correlation of the true grouping scores with the grouping scores from the standard hallmark structure matrices and the meta-structure matrix. The left column shows the correlations under the strong-anchor setting; the patterns are very similar to those of the prioritization score correlations. As with the prioritization scores, there is a significant penalty for overspecifying the number of topics under the standard method, but SEEK-VEC, which includes these overspecified models, does not experience the same drop in performance. The right column shows the correlations as the anchor condition weakens. Again, SEEK-VEC outperforms the standard models, demonstrating the value of the ensemble method under weaker signal strength conditions.

#### SEEK-VEC provides a lens for model diagnostics

The output matrix from SEEK-VEC can also be used to assess the stability of a given topic model. Comparing the hallmark structure matrix from a standard topic model to the SEEK-VEC output can inform the extent to which the conclusions of the proposed topic model are conditional on the choice of *K*. For example, suppose 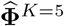 is the hallmark structure matrix corresponding to a standard topic model with *K* = 5 topics and 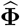 is the meta-structure matrix from SEEK-VEC. A high value of 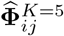 and low value of 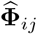 indicates that the signal from the *K* = 5 model may be unique to that particular model and therefore should be interpreted with caution.

Fig. 3 shows heatmaps representing standard topic models for *K* = {3, …, 8} based on data generated under a true *K* = 6 model. Each heatmap includes top 10 words for each derived topic, and the heatmap is shaded by the corresponding value from the SEEK-VEC matrix. The top left heatmap, which shows the top words from the *K* = 3 model, indicates that the topic represented by the middle block is slightly less stable across models than the other two topics: the lighter colors indicate that there are some models for which some of the words in the first topic are not identified as representative words for the same topics. Similarly, the *K* = 4 model shows that the top words in the second topic are more consistently co-identified as representative words than the top words for the other topics, and can thus be seen as more stable with respect to hyperparameter selection.

**Figure 3:**
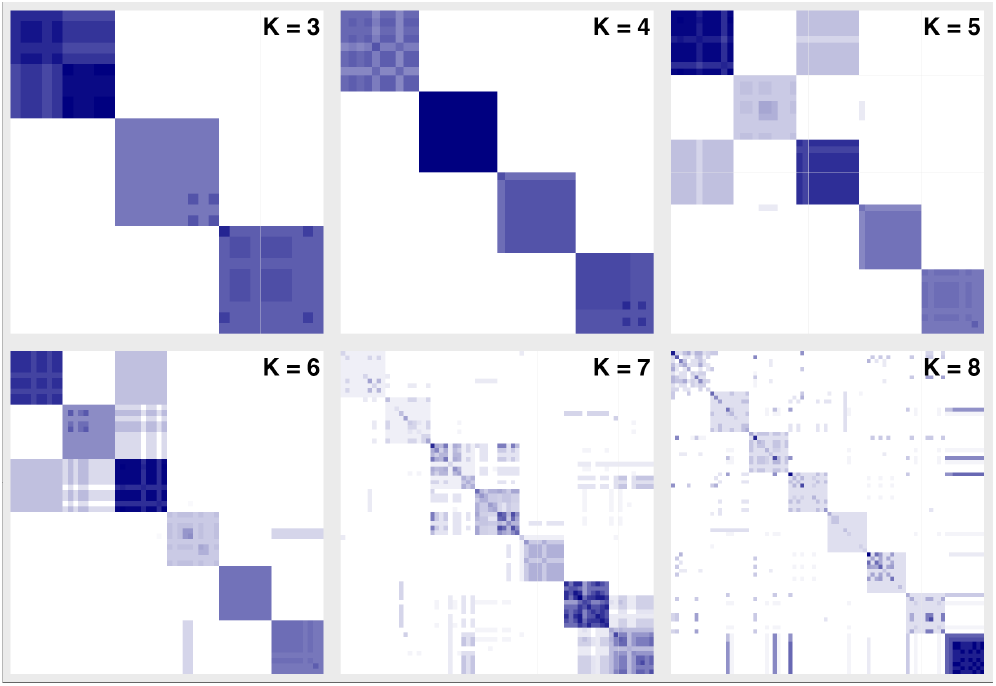
Heatmaps (via *pheatmap*^23^) showing the top 10 words in each component topic model, colored by their value in the SEEK-VEC meta-structure matrix.

The *K* = 5 model demonstrates some interrelationships among the five topics. The shaded areas on the off-diagonal indicate that there are some models for which words in the first and third topics in the *K* = 5 model are co-identified as representative words. In other words, in other candidate models, while words in these two blocks are hallmark for separate topics in the *K* = 5 model, in other models they share a topic. This is shown further in the output from the correctly-specified *K* = 6 model. This plot shows a relationship between the first and third topics, as well as the to a lesser degree the fourth and sixth topics. In the case of randomly-simulated data, any meta-relationships among the topics are incidental; however, in the real data case, this can indicate topics that are more closely related. For example, if two topics are “baseball” and “‘basketball”, these topics may have been agglomerated into a “sports” topic in an underspecified model, and therefore would show a relationship along the off-diagonal in the ensemble matrix.

Finally, the last two heatmaps correspond to the overspecified *K* = 7 and *K* = 8. As seen in Supplemental Fig. A5, overspecifying *K* leads to noisy estimation of the topics. This is shown clearly in these figures by the lack of consistently-strong block structure along the diagonal. Blocks that are barely shaded represent topics that are potentially unique to that specific topic model, and are therefore very unstable with respect to the selection of *K*. Interpretation of these topics should be done with caution, as they may not represent true signal but rather noisy artifacts of the choice of hyperparameter. A real data example of this is found in Section 2.3.

#### SEEK-VEC is more robust to hyperparameter selection

There are two hyperparameters required for SEEK-VEC: the choice of values of *K* to ensemble and the threshold value *t*. Fig. 2c shows the performance of SEEK-VEC ensembling *K* = {3, …, *K*\*} across values of *K*\*. Across both anchor strength settings and both sets of scores, the pattern is consistent: all models with *K*\* ≥ 6 have near-identical performance. Compared to the performance of the standard topic models in Fig. 2b, this shows that there is much less risk in choosing *K*\* for SEEK-VEC than in choosing *K* for Topic-SCORE. In particular, for SEEK-VEC, there is no penalty in overestimation, while there is a significant penalty for Topic-SCORE. Fig. 2d and Fig. 2e show direct comparisons of Topic-Score with *K* = *k* versus SEEK-VEC with *K*\* = *k* for prioritization scores and grouping scores, respectively. With the exception of *k* = 6 in the strong anchor condition, when Topic-SCORE is an oracle method, there is either no harm or an improvement in performance for implementing SEEK-VEC. Supplemental Fig. A4 shows results equivalent to Fig. 2c for the ensemble strategies of uniform averaging and Frobenius normalized averaging and demonstrates that these methods are much less robust to hyperparameter selection compared to SEEK-VEC.

As SEEK-VEC is intended to complement a standard topic modeling procedure, the appropriate choice of *t* is that which replicates what is chosen for the standard procedure.

#### SEEK-VEC requires minimal additional computational time

Fig. 2f shows the additional computational time required by the ensembling step compared to the computational time required by running the component topic models across a variety of topic modeling methods. The rows represent different topic modeling algorithms, and the columns represent different values of the vocabulary size *p*. For all methods except for Topic-SCORE, the added computational time to run the ensembling step is trivial relative to the time required to run the standard models. For Topic-SCORE, for *p* ≥ 1000, the ensembling step using a higher threshold value *t* can take up to twice as long as running the component models. As shown in Algorithm 3, the input to the spectral ensembling procedure is a set of *p*\* × *p*\* matrices, where *p*\* ≤ *p* is the number of vocabulary words with non-zero diagonal entries across all component matrices. Therefore, choosing a lower value of *t* means a smaller value of *p*\*, which means smaller input matrices for the spectral ensembling step. However, the absolute additional time required when using Topic-SCORE to ensemble *K* = {3, …, 12}, even with larger *p* and higher threshold value *t*, is at most just over a minute.

### 2.3 Application of SEEK-VEC to the MADStat Dataset

The MADStat dataset is a dataset compiled by the authors of Topic-SCORE of the abstracts of 83,331 research papers published in 36 statistical journals between 1975 and 2015^1^. Based on the scree plot of **D**, they identified *K* = {4, …, 16} as a reasonable range of values to consider. For each candidate *K*, the authors ran Topic-SCORE, obtained a topic loadings matrix, assessed the interpretability of the top 20 words per topic and chose *K* = 11 based on subject-matter expertise. The authors advocate for this subjective procedure for selecting *K*, stating that data-driven methods can be unstable or biased towards uninterpretable models with large values of *K*. Here we apply SEEK-VEC to the MADStat data. To directly compare our results to that of the original authors, we ensemble models *K* = {4, …, 16}. We first dichotomize the resulting topic loadings matrices by choosing the top 20 words per topic, to match the original procedure 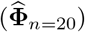. We also dichotomize using the threshold method to assess the overall distribution of probabilities across the word-topic space 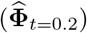.

First, we can examine the authors’ chosen *K* = 11 model with respect to the ensemble method. Fig. 4a shows a heatmap of the top 20 representative words per topic from the *K* = 11 model. The heatmap is colored by 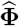 using the dichotomization strategy of identifying the 20 words with the highest loadings per topic 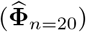, which simulates how the MADStat authors evaluated the candidate topic models. The block corresponding to *Exp Design* is heavily shaded, indicating that these words are frequently co-identified as representative words. This topic, therefore, is stable with respect to the choice of the hyperparameter *K*. The *Bio/Med* topic, on the other hand, is much lighter, indicating that this topic is more sensitive to hyperparameter selection. Fig. 4b again displays the top 20 most representative words per topic, but this time colored by 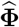 using the threshold strategy with *t* = 0.2 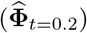. This method does not enforce equal representation of topics and therefore gives a more accurate representation of signal strength within the data. Here, we can see a strong interrelationship between the *Bio/Med* topic and the *Clinic* topic, which is reasonable given that these two topics both fall under the category of biostatistics. We can also see an interrelationship between these two topics and the *Time Series* topic. As before, the *Exp Design* shows high stability, with heavy weights within the topic and little to no overlap with other topics.

**Figure 4:**
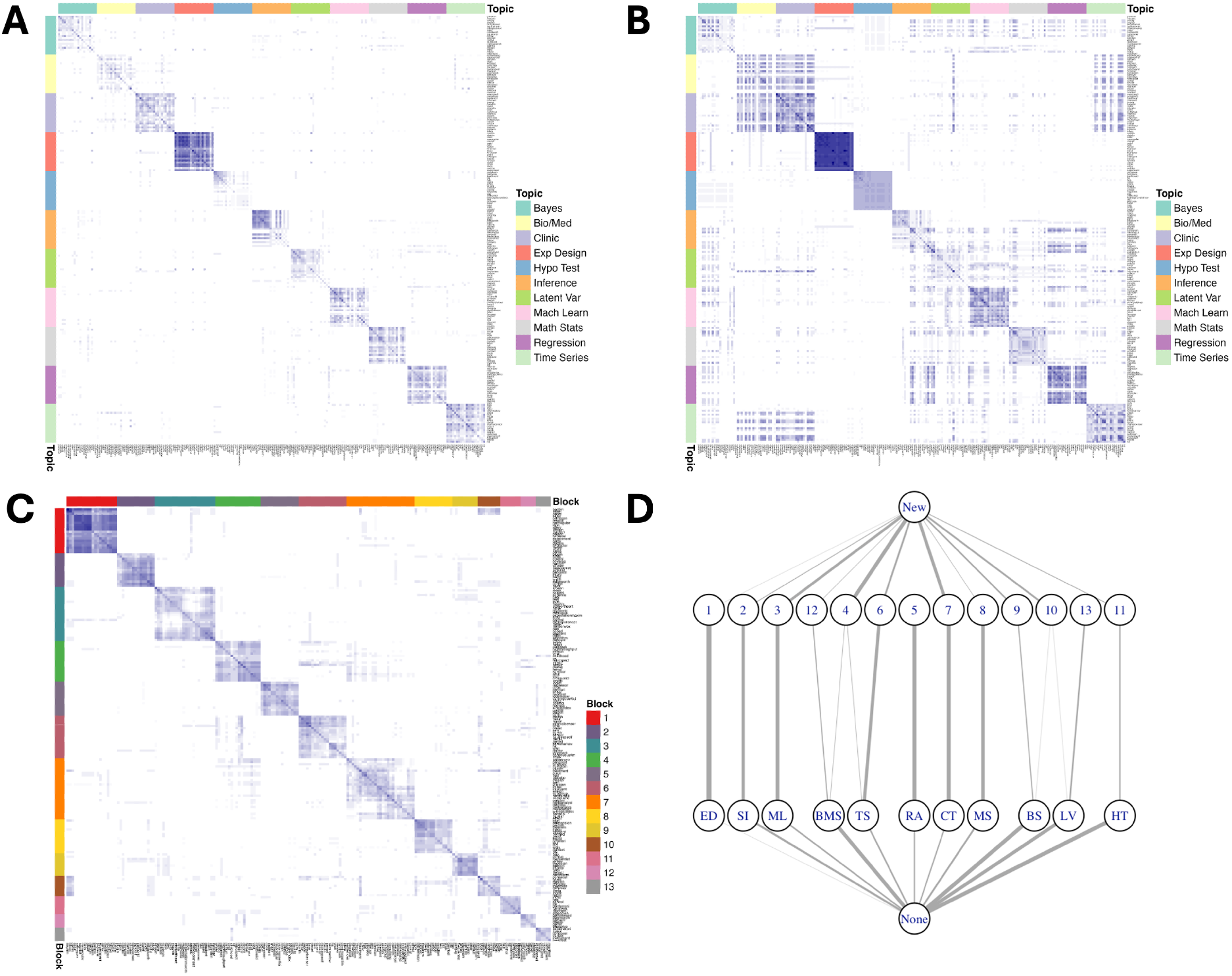
(a-c) Heatmaps of representative words and researcher-derived topics corresponding to the *K* = 11 model in the original MADStat paper. (a) All 20 representative words used for topic labeling colored by 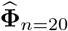, the meta-structure matrix derived using the dichotomization strategy of choosing the top 20 words from each topic (simulating the strategy used in the MADStat paper). (b) The 20 words with the strongest loadings for each topic colored by 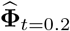, the ensemble matrix derived using the dichotomization strategy of thresholding by the top 20% of strongest topic loadings. (c) The topic blocks derived from 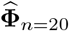. The heatmap is filtered to the top 10% of words by diagonal value. (d) Graph showing the relationship between the *K* = 11 topics and the SEEK-VEC topic blocks derived from 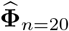. The “New” node represents words highlighted by the ensemble method that were not representative words in the *K* = 11 method, and “None” represents words that were identified as representative by the *K* = 11 method but not by the ensemble method.

Fig. 4c shows the block structure that emerges when restricting 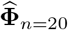 to the top 10% of words as by diagonal value, and Fig. 4d shows the relationship between these blocks and the representative words selected by the *K* = 11 model. The lists of words per block from 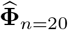 are shown in Section A.2.8. Many of the blocks strongly correspond to topics from the *K* = 11 model. For example, Block 1 has a strong relationship to the *Exp Design* topic: most words are identified by both models, with only one word identified in the *Exp Design* topic and not Block 1 (“column”), and only one word identified in Block 1 and not the *Exp Design* topic (“fraction”). Some of the blocks represent combinations of topics from the *K* = 11 model, such as Block 4, which draws on both *Bio/Med* and *Time Series*. Block 5 is a subset of *Regression* and Block 13 is a subset of *Latent Var*, although *Latent Var* also shares words with Block 10, which also has some overlap with *Bayes* primarily comprises newly-identified words.

In this section, we demonstrated the utility of SEEK-VEC to evaluate the robustness of a selected topic model across candidate values of *K* and provided an alternative strategy for identifying important vocabulary words by directly working with the meta-structure matrix. A more exploratory use case of SEEK-VEC in a non-text setting is found in Section A.2.6.

## 3 Discussion

Topic modeling is a powerful tool for detecting latent structure in count data. However, standard topic models require choosing a number of topics *K*, which may not be possible in non-text data settings, such as symptom data or gene expression data, either due to a lack of interpretability of the candidate topic models or weaker signal strength in the absence of the anchor word condition. SEEK-VEC is a framework for augmenting topic models that is much less sensitive to hyperparameter selection and improves performance in weaker signal strength scenarios. As shown through simulations, SEEK-VEC can improve the identification of topic-relevant vocabulary words and stable relationships between words. When applied to the MADStat data, SEEK-VEC yields insights into the stability of a given model output, feature selection across models, and alternative possible grouping structure within the data.

A key limitation of SEEK-VEC is that it requires the construction and analysis of *p* × *p* matrices. It is therefore best suited for settings with small-to-moderate *p*, including high-dimensional settings where *p* << *n*, such as surveys with many more respondents than items.Future development of SEEK-VEC to extend to large data contexts would necessitate efficiency improvements along the line of the Zero-Reduce step currently in the algorithm; for example, if each candidate 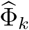 comprises several disconnected components, the ensemble approach could be done in a piece-wise manner. Further, replacing the matrix multiplication step with a parallel algorithm would likewise make the method more efficient. Another area of future research to make SEEK-VEC more suitable for large-*p* contexts is to instead ensemble the latent subspace in which 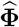 can be thought of as a distance matrix.

Another avenue for future exploration with SEEK-VEC is its utility for understanding the behavior of black-box algorithms with respect to their hyperparameters. AI-based methods such as BERTopic^24^ lack a statistical framework and can be seen as black-box methods, meaning it can be challenging to assess the stability and reliability of the output.^25,26^ Future work with SEEK-VEC could explore its use as a non-parametric tool to analyze the stability of the output of black-box models with respect to their hyperparameters without requiring any statistical framework for the method itself. A preliminary example of this use case, which evaluates the stability of BERTopic with respect to the underlying random seed, is found in Supplemental Section A.2.7.

## Acknowledgements

R.D.’s work was supported by NIH Grant T32-GM135117. Z.T.K.’s work is supported by the Sloan Research Grant FG-2023-19970. X.L.’s work is supported by NIH Grants R35-CA197449, R01-HL163560, and U01-HG012064.

## References

1. Ke, Z. T., Ji, P., Jin, J. & Li, W. Recent Advances in Text Analysis. Annual Review of Statistics and Its Application 11, 347–372. ISSN: 2326-831X. 10.1146/annurev-statistics-040522-022138 (Apr. 2024).

2. Bicego, M., Lovato, P., Ferrarini, A. & Delledonne, M. Biclustering of Expression Microarray Data with Topic Models in 2010 20th International Conference on Pattern Recognition (IEEE, Aug. 2010), 2728–2731. 10.1109/ICPR.2010.668.

3. Konietzny, S. G., Dietz, L. & McHardy, A. C. Inferring functional modules of protein families with probabilistic topic models. BMC Bioinformatics 12. ISSN: 1471-2105. 10.1186/1471-2105-12-141 (May 2011).

4. Castellani, U. et al. in Medical Image Computing and Computer-Assisted Intervention – MICCAI 2010 177–184 (Springer Berlin Heidelberg, 2010). ISBN: 9783642157455. 10.1007/978-3-642-15745-5_22.

5. Liu, L., Tang, L., Dong, W., Yao, S. & Zhou, W. An overview of topic modeling and its current applications in bioinformatics. SpringerPlus 5. ISSN: 2193-1801. 10.1186/s40064-016-3252-8 (Sept. 2016).

6. Fan, A., Doshi-Velez, F. & Miratrix, L. Assessing topic model relevance: Evaluation and informative priors. Statistical Analysis and Data Mining: The ASA Data Science Journal 12, 210–222. ISSN: 1932-1872. 10.1002/sam.11415 (May 2019).

7. Ke, Z. T. & Wang, M. Using SVD for Topic Modeling. Journal of the American Statistical Association 119, 434–449. ISSN: 1537-274X. 10.1080/01621459.2022.2123813 (Oct. 2022).

8. Blei, D. M., Ng, A. Y. & Jordan, M. I. Latent Dirichlet Allocation. J. Mach. Learn. Res. 3, 993–1022. ISSN: 1532-4435 (Mar. 2003).

9. Arora, S. et al. A Practical Algorithm for Topic Modeling with Provable Guarantees in Proceedings of the 30th International Conference on Machine Learning (eds Dasgupta, S. & McAllester, D.) 28 (June 2013), 280–288.

10. Bansal, T., Bhattacharyya, C. & Kannan, R. A provable SVD-based algorithm for learning topics in dominant admixture corpus in Neural Information Processing Systems (2014). https://api.semanticscholar.org/CorpusID:15386282.

11. Fu, Q., Zhuang, Y., Gu, J., Zhu, Y. & Guo, X. Agreeing to Disagree: Choosing Among Eight Topic-Modeling Methods. Big Data Research 23, 100173. ISSN: 2214-5796. 10.1016/j.bdr.2020.100173 (Feb. 2021).

12. Zhao, W. et al. A heuristic approach to determine an appropriate number of topics in topic modeling. BMC Bioinformatics 16. ISSN: 1471-2105. 10.1186/1471-2105-16-S13-S8 (Dec. 2015).

13. Wang, H. & Miller, D. Improved Parsimonious Topic Modeling Based on the Bayesian Information Criterion. Entropy 22, 326. ISSN: 1099-4300. 10.3390/e22030326 (Mar. 2020).

14. Arora, S., Ge, R. & Moitra, A. Learning Topic Models – Going beyond SVD in 2012 IEEE 53rd Annual Symposium on Foundations of Computer Science (IEEE, Oct. 2012), 1–10. 10.1109/FOCS.2012.49.

15. Donoho, D. & Stodden, V. When does non-negative matrix factorization give a correct decomposition into parts? Proceedings of the 17th International Conference on Neural Information Processing Systems 16, 1141–1148 (2003).

16. Ding, W., Ishwar, P. & Saligrama, V. Most large topic models are approximately separable in 2015 Information Theory and Applications Workshop (ITA) (IEEE, Feb. 2015), 199–203. 10.1109/ITA.2015.7308989.

17. Freyaldenhoven, S., Ke, S., Li, D. & Montiel Olea, J. L. On the Testability of the Anchor-Words Assumption in Topic Models. Working paper (Federal Reserve Bank of Philadelphia). ISSN: 2574-0997. 10.21799/frbp.wp.2025.14. (Apr. 2025).

18. Kittler, J., Hatef, M., Duin, R. & Matas, J. On combining classifiers. IEEE Transactions on Pattern Analysis and Machine Intelligence 20, 226–239. ISSN: 0162-8828. 10.1109/34.667881 (Mar. 1998).

19. He, H. & Cao, Y. SSC: A Classifier Combination Method Based on Signal Strength. IEEE Transactions on Neural Networks and Learning Systems 23, 1100–1117. ISSN: 2162-2388. 10.1109/TNNLS.2012.2198227 (July 2012).

20. Banerjee, S., Jenamani, M. & Pratihar, D. K. Properties of a projected network of a bipartite network in 2017 International Conference on Communication and Signal Processing (ICCSP) (IEEE, Apr. 2017), 0143–0147. 10.1109/ICCSP.2017.8286734.

21. Ma, R., Sun, E. D. & Zou, J. A spectral method for assessing and combining multiple data visualizations. Nature Communications 14. ISSN: 2041-1723. 10.1038/s41467-023-36492-2 (Feb. 2023).

22. Zipf, G. K. The Psycho-Biology Of Language: An Introduction to Dynamic Philology (Houghton Mifflin, Boston, 1935).

23. Kolde, R. pheatmap: Pretty Heatmaps R package version 1.0.12 (2019). https://CRAN.R-project.org/package=pheatmap.

24. Grootendorst, M. BERTopic: Neural topic modeling with a class-based TF-IDF procedure. arXiv preprint arXiv:2203.05794 (2022).

25. Raman, R., Pattnaik, D., Hughes, L. & Nedungadi, P. Unveiling the dynamics of AI applications: A review of reviews using scientometrics and BERTopic modeling. Journal of Innovation & Knowledge 9, 100517. ISSN: 2444-569X. 10.1016/j.jik.2024.100517 (July 2024).

26. Sadeghi, Z. et al. A review of Explainable Artificial Intelligence in healthcare. Computers and Electrical Engineering 118, 109370. ISSN: 0045-7906. 10.1016/j.compeleceng.2024.109370. (Aug. 2024).

27. Parisi, F., Strino, F., Nadler, B. & Kluger, Y. Ranking and combining multiple predictors without labeled data. Proceedings of the National Academy of Sciences 111, 1253–1258. ISSN: 1091-6490. 10.1073/pnas.1219097111 (Jan. 2014).

28. Pedregosa, F. et al. Scikit-learn: Machine Learning in Python. Journal of Machine Learning Research 12, 2825–2830 (2011).

29. Blei, D. & Lafferty, J. Correlated Topic Models in NIPS’05: Proceedings of the 19th International Conference on Neural Information Processing Systems (Dec. 2005), 147–154.

30. Carbonetto, P., Sarkar, A., Wang, Z. & Stephens, M. Non-negative matrix factorization algorithms greatly improve topic model fits 2021. https://arxiv.org/abs/2105.13440.

31. Teh, Y., Jordan, M., Beal, M. & Blei, D. Sharing Clusters among Related Groups: Hierarchical Dirichlet Processes in Advances in Neural Information Processing Systems (eds Saul, L., Weiss, Y. & Bottou, L.) 17 (MIT Press, 2004). https://proceedings.neurips.cc/paper_files/paper/2004/file/fb4ab556bc42d6f0ee0f9e24ec4d1af0-Paper.pdf.

32. Wallach, H. M., Murray, I., Salakhutdinov, R. & Mimno, D. Evaluation methods for topic models in Proceedings of the 26th Annual International Conference on Machine Learning (ACM, 2009), 1105–1112. 10.1145/1553374.1553515.

33. Wang, C., Paisley, J. & Blei, D. M. Online Variational Inference for the Hierarchical Dirichlet Process in Proceedings of the Fourteenth International Conference on Artificial Intelligence and Statistics (eds Gordon, G., Dunson, D. & Dudík, M.) 15 (PMLR, Fort Lauderdale, FL, USA, Nov. 2011), 752–760. https://proceedings.mlr.press/v15/wang11a.html.

34. Teh, Y. W., Jordan, M. I., Beal, M. J. & Blei, D. M. Hierarchical Dirichlet Processes. Journal of the American Statistical Association 101, 1566–1581. ISSN: 1537-274X. 10.1198/016214506000000302 (Dec. 2006).

35. Simons Foundation Autism Research Initiative. About SPARK Accessed: 2025-02-03. Sept. 2024. https://sparkforautism.org/portal/page/about-spark/.

36. De Vries, L. P. et al. A Comparison of the ASEBA Adult Self Report (ASR) and the Brief Problem Monitor (BPM/18-59). Behavior Genetics 50, 363–373. ISSN: 1573-3297. 10.1007/s10519-020-10001-3 (May 2020).

37. Achenbach, T. M. DSM Guide for the ASEBA (University of Vermont, Research Center for Children, Youth, & Families, Burlington, VT, 2013).

38. Xi, T. & Wu, J. A Review on the Mechanism Between Different Factors and the Occurrence of Autism and ADHD. Psychology Research and Behavior Management Volume 14, 393–403. ISSN: 1179-1578. 10.2147/PRBM.S304450 (Apr. 2021).

39. Storebø, O. J. & Simonsen, E. The Association Between ADHD and Antisocial Personality Disorder (ASPD): A Review. Journal of Attention Disorders 20, 815–824. ISSN: 1557-1246. 10.1177/1087054713512150 (July 2016).

40. Hudson, C. C., Hall, L. & Harkness, K. L. Prevalence of Depressive Disorders in Individuals with Autism Spectrum Disorder: a Meta-Analysis. Journal of Abnormal Child Psychology 47, 165–175. ISSN: 1573-2835. 10.1007/s10802-018-0402-1 (Mar. 2018).

41. Hinze, E., Paynter, J., Dargue, N. & Adams, D. The Presentation of Depression in Depressed Autistic Individuals: A Systematic Review. Review Journal of Autism and Developmental Disorders. ISSN: 2195-7185. 10.1007/s40489-024-00480-z (Sept. 2024).

42. Rodgers, J. & Ofield, A. Understanding, Recognising and Treating Co-occurring Anxiety in Autism. Current Developmental Disorders Reports 5, 58–64. ISSN: 2196-2987. 10.1007/s40474-018-0132-7 (Jan. 2018).

43. McInnes, L., Healy, J. & Melville, J. UMAP: Uniform Manifold Approximation and Projection for Dimension Reduction 2018. https://arxiv.org/abs/1802.03426.

