## Supplemental materials for "SEEK-VEC: Augmenting topic modeling with spectral ensemble learning"

### A Appendix

#### A.1 Methods and Materials

##### A.1.1 Topic Modeling Framework

Given $n$ observed documents written on a vocabulary of $p$ words, Let $\mathbf{D} \in \mathbb{R}^{p \times n}$ be a matrix comprising the counts of each vocabulary word in each document. Let $\mathbf{A} \in \mathbb{R}^{p \times K}$ be a latent matrix representing each vocabulary word's weight on each of $K$ topics, such that $A_1, ..., A_K$, the columns of $\mathbf{A}$, represent probability mass functions on the vocabulary. A word in the vocabulary is defined as an *anchor word* if its corresponding row in the $\mathbf{A}$ matrix has only one non-zero entry; these words can be thought to identify the topics. Let $\mathbf{W} \in \mathbb{R}^{K \times n}$ be another latent matrix which represents the mixture of topics within each document, with $w_i(k)$ representing the weight of document $i$ on topic $k$. For a document $i$ of length $N_i$, topic modeling posits that the $i$-th column of $\mathbf{D}$ is drawn as $N_i d_i \sim Multinomial(N_i, \sum_{k=1}^{K} w_i(k) A_k)$. This yields the following relationship:

$$\mathbb{E}[\mathbf{D}] = \mathbf{AW}$$

Since typically $K << n$, this implies a low-rank structure underlying $\mathbf{D}$[7].

##### A.1.2 Topic-SCORE

Topic-SCORE is a state-of-the-art method for topic modeling that uses singular value decomposition and simplex geometry to estimate the low-rank $\mathbf{A}$ from $\mathbf{D}$[7]. After initial normalization of the observed $\mathbf{D}$ matrix to account for word frequency heterogeneity, Topic-SCORE obtains the matrix of the $K$ left singular vectors $\Xi = [\xi_1 ... \xi_K]$. The rows of $\Xi$ are contained in a simplicial cone with $K$ supporting rays, where the anchor words lie on the supporting rays. Next, to compress the simplicial cone into a simplex, $\Xi$ is normalized via SCORE normalization: each row is normalized by its first component, yielding $\mathbf{R} \in \mathbb{R}^{p \times (K-1)}$. Each row of $\mathbf{R}$ represents a low-dimensional embedding of a word into $\mathbb{R}^{K-1}$. The anchor words, which were located on the supporting rays of the $K$-dimensional simplicial cone, are now located at the vertices of the $(K-1)$-dimensional simplex. These vertices are found using a vertex hunting step, and since these vertices identify the location of the pure topics, each vocabulary word's weight on each topic can be found by uncovering the convex combination of vertices that yields its position in the simplex, thus yielding the desired low-rank matrix $\mathbf{A} \in \mathbb{R}^{p \times K}$. Importantly, misspecifying $K$ implies the embedding of the vocabulary onto a space with the incorrect number of dimensions; the impact of this misspecification is further explored in Fig. A5.

##### A.1.3 Details of SEEK-VEC

The target estimand in topic modeling is $\mathbf{A}$, a $p \times K$ matrix representing the weight of each of $p$ words on each of $K$ topics. Each column of $\mathbf{A}$ represents a probability mass function (PMF) on the vocabulary, with the most frequently-used words per topic having the highest probability. This means that the rows of $\mathbf{A}$ corresponding to common words such as "and" or "the" will have high probabilities in each column, despite not being useful for the identification or interpretation of the topics. Thus, in practice, what is usually used for the interpretation of the topics is a row-normalized version of $\mathbf{A}$ called the topic loadings matrix[1]. Each row of the topic loadings matrix represents the relative weights of each topic for each word.

The "representative words" for each topic are found by identifying the words in the topic loadings matrix with the largest values[7]. This can be done in several ways, such as choosing the top $n$ words per column or by choosing the $n$ words with the largest values across the entire topic loadings matrix. The advantage of the former strategy is that it ensures the same number of representative words per topic and prevents a topic with many strong representative words from dominating the interpretation; the advantage for the latter strategy is it avoids inflating the importance of relatively unimportant words in the case where a given topic has fewer truly informative words than others. We can represent this differentiation process between representative words and uninformative words as a dichotomizing of the topic loadings matrix to the hallmark matrix $\mathbf{H}$. In this matrix, $h_{ij} = 1$ if word $i$ is representative for topic $j$, and 0 otherwise. This yields the hallmark structure matrix $\mathbf{\Phi} = \mathbf{HH}^\top$.

In order to understand the topic structure with respect to the vocabulary without knowing $K$, we can estimate $\mathbf{\Phi} \in \mathbb{R}^{p \times p}$ by combining the insights of a variety of candidate topic models. To develop an estimator of $\mathbf{\Phi}$, we propose the following spectral ensembling method:

Let $\mathcal{K} = \{3, ..., K^*\}$ be a collection of candidate topic counts. For each $k \in \mathcal{K}$, let $\widehat{\mathbf{A}}_k$ be the topic matrix derived from running Topic-SCORE with $k$ topics. Let $\widehat{\mathbf{\Phi}}_k$ be the corresponding hallmark structure matrix.

Define the similarity matrix $\mathbf{G} \in \mathbb{R}^{|\mathcal{K}| \times |\mathcal{K}|}$ where

$$\mathbf{G}(i, j) = tr((\widehat{\mathbf{\Phi}}_{k_i})^\top \widehat{\mathbf{\Phi}}_{k_j}),$$

which measures the similarity between the hallmark structure matrix $\widehat{\mathbf{\Phi}}_{k_i}$ using $k_i$ topics and the hallmark structure matrix $\widehat{\mathbf{\Phi}}_{k_j}$ using $k_j$ topics. Let $u_1 \in \mathbb{R}^{|\mathcal{K}|}$ be the leading eigenvector of $\mathbf{G}$. Then we can combine the $|\mathcal{K}|$ hallmark structure matrices as follows:

$$\widehat{\mathbf{\Phi}} = \sum_{k \in \mathcal{K}} \frac{|u_{1k}|}{||u_1||_1} \widehat{\mathbf{\Phi}}_k$$

The resulting matrix $\widehat{\mathbf{\Phi}}$ is a consensus-weighted combination of the hallmark structure matrices from the candidate topic models (Algorithm 3). Since $\widehat{\mathbf{\Phi}}$ is a weighted combination of integer-valued matrices, it will not itself necessarily (or likely) be integer-valued. However, as is shown, it can still be used as a guide to identifying topic-relevant vocabulary words and labeling the topics.

This spectral ensembling method generalizes the spectral meta-learner method of Parisi et al. (2014)[27] and the eigenscore method of Ma et al. (2023)[21]. Both methods address the question of the best way to combine candidate models via a consensus-based weighting, where the leading eigenvector of a matrix summarizing the differences among the models provides an optimal weighting under modest regularity assumptions. The spectral meta-learner method assumes conditional independence across component models[27], which may be violated in the topic modeling setting because the topics from candidate models can be mixtures of the oracle model[7]. However, the eigenscore method does not assume complete independence and allows for weak to moderate correlation among the input matrices[21]. The default topic modeling method used in this paper is Topic-SCORE; however, alternate methods can be used if desired (see Section A.2.1).

###### A.1.4 Method Implementation

The overview of the implementation of SEEK-VEC is given in Algorithm 1. As with standard topic modeling methods, the method requires the word-document corpus matrix $\mathbf{D}$ as input. The method also requires $\mathcal{K}$, which is a list of candidate topic counts to ensemble. Finally, the method requires $t$, the parameter for thresholding the topic loadings matrices. Details of this procedure are shown in Algorithm 2, which involves converting each output $\mathbf{A}$ matrix to a topic loadings matrix by normalizing the rows to sum to 1. The loadings matrix is then converted to a binary matrix through the thresholding procedure, which is then transpose-multiplied by itself to obtain a $p \times p$ matrix that is free of $k$ and can thus be ensembled through the spectral ensembling procedure. This procedure, shown in more detail in Algorithm 3, is optimized for computational efficiency through a Zero-Reduce procedure. For this step, any vocabulary word which has a diagonal value of 0 across all candidate matrices is removed prior to calculating the $\mathbf{G} \in \mathbb{R}^{|\mathcal{K}| \times |\mathcal{K}|}$ matrix, since any weighted sum of the candidate matrices will result in zeroes across the rows and columns of these vocabulary words. Once $\mathbf{G}$ is calculated, the candidate matrices are combined in a weighted sum proportional to the leading eigenvector of $\mathbf{G}$. The output of the algorithm is the meta-structure matrix $\Phi$. The prioritization score for the $i^{th}$ vocabulary word is $\Phi_{ii}$, and the grouping score for the $i^{th}$ and $j^{th}$ vocabulary words is $\Phi_{ij} = \Phi_{ji}$.

**Algorithm 1** SEEK-VEC

**Input:** $D \in \mathbb{R}^{p \times n}, \mathcal{K}, t$

  **for** $k \in \mathcal{K}$ **do**

    $\widehat{A}_k \leftarrow$ Topic-SCORE$(D, k)$

    $\widehat{\Phi}_k \leftarrow$ A-to-$\Phi(\widehat{A}_k, t)$

  $\widehat{\Phi} \leftarrow$ Spectral-Ensemble$(\{\widehat{\Phi}_{k \in \mathcal{K}}\})$

**Output:** $\widehat{\Phi}$

**Algorithm 2** A-to-$\Phi$

**Input:** $A \in \mathbb{R}^{p \times k}, t$

  $B \leftarrow$ Row-Normalize$(A)$

  $H \leftarrow$ Threshold$(B, t)$

  $\Phi \leftarrow HH^\top$

**Output:** $\Phi$

**Algorithm 3** Spectral-Ensemble

**Input:** $\Phi_1, ..., \Phi_n \in \mathbb{R}^{p \times p}$

  $\Phi_1^*, ..., \Phi_n^* \leftarrow$ Zero-Reduce$(\Phi_1, ..., \Phi_n)$

  $G \in \mathbb{R}^{n \times n}$

  $G_{i,j} \leftarrow tr({\Phi^*}_i^\top {\Phi^*}_j)$

  $u_1 \in \mathbb{R}^n \leftarrow$ Leading-Eigenvector$(G)$

  $\Phi \leftarrow \sum_{i=1}^n \frac{|u_{1i}|}{||u_1||_1} O_i$

**Output:** $\Phi$

##### A.1.5 Data and Code Availability

SEEK-VEC is available to download as an R package at `https://github.com/rdanning/seekvec`. The code and parameters for generating the simulation data and replicating the real data analysis is available at `https://github.com/rdanning/SEEK-VEC-repro`. The MADStat data is available to download from GitHub at https://github.com/ZhengTracyKe/MADStat. The SPARK Adult Self-Report is available through application to the Simons Foundation (https://www.sfari.org/resource/spark/). The 20 newsgroups dataset is available through the scikit-learn package.[28]

#### A.2 Supplementary Materials

##### A.2.1 Choosing a topic modeling method for SEEK-VEC

There are several methods that can be used in conjunction with the SEEK-VEC framework to estimate the underlying word-topic matrix **A**. In this work, we focus on four methods that have software implementations in the *R* programming language. A brief discussion of alternate methods is found in Section A.2.2. Latent Dirichlet Allocation (LDA)[8] is a Bayesian mixture model that assumes the vocabulary are generated by a hierarchical process based on the proportions of orthogonal topics in each document. Correlated Topic Models (CTM)[29] is similar to LDA but expands the method to allow for correlations among the topics. Another method, fastTopics, uses Poisson non-negative matrix factorization to estimate **A**[30]. Topic-SCORE[7] is a recent approach based on singular value decomposition and simplex geometry that directly addresses the word frequency heterogeneity commonly found in text data. As shown in Supplemental Fig. A1a, all methods are able to accurately estimate the diagonal of the meta-structure matrix $\mathbf{\Phi}$ when the true $K$ is known and there is strong signal in the data, i.e. the anchor word condition holds strongly.

**Ensembling improves performance across methods and hypothesized numbers of topics** In the case when $K$ is not known, Supplemental Fig. A1b shows the change in performance comparing the standard implementation of the method (e.g. running the method with $K = K^*$) to the ensemble method (e.g. ensembling $K = \{3, ..., K^*\}$) for varying values of $K^*$. For LDA, fastTopics, and Topic-SCORE, it is always significantly advantageous to ensemble in the case that the number of topics has been overspecified. For LDA and fastTopics, there is no significant difference with ensembling when the number of topics is either correctly- or under-specified. For Topic-SCORE, there is no significant difference for ensembling when the number of topics is underspecified, and only a slight penalty for ensembling when the number of topics is correctly specified. For CTM, the results are only statistically significant at higher values of $K^*$, but there is no significant penalty at any value for ensembling.

However, while all methods perform well under strong signal, this is not the case under weaker signal conditions, which often occur in non-text practical data settings, such as single cell expression or survey data. Supplemental Fig. A1c shows the diagonal correlation results comparing five methods under the setting when the true $K = 6$ is known. The anchor words increase in strength along the $x$-axis, so the noisier settings correspond to the left side of the plots. Only fastTopics and Topic-SCORE are robust to settings where the anchor word condition is moderately violated.

(a) Results with true $K$ (b) Ensembling improvement (c) Performance with noise

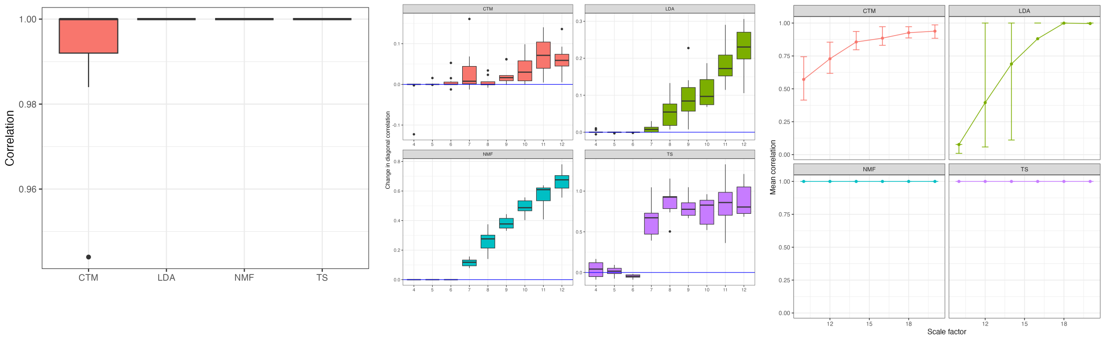


Supplemental Figure A1: Comparisons of the four main topic modeling methods: Correlated Topic Models (“CTM"), Latent Dirichilet Allocation (“LDA"), fastTopics (“NMF"), and TopicScore (“TS"). The vocabulary size is $p = 500$, the corpus size is $n = 1,000$ documents, and there are $N = 2,000$ words per document. There are no metainformative words, 50% of words are identifying words, and we do not enforce word frequency heterogeneity. The true number of topics is $K = 6$. (a,b) True anchor condition. (c)The scale factor represents the ratio between loadings for representative and non-representative words for a topic; as the ratio increases, the representative words become more like true anchor words. Error bars represent the $25^{th}$ and $75^{th}$ percentiles.

**Topic-SCORE is much more efficient than fastTopics** As shown in Supplemental Fig. A2, when the vocabulary size, number of documents, and true number of topics increases, Topic-SCORE becomes much more efficient than fastTopics. As such, we focus on implementing Topic-SCORE in this paper.

##### A.2.2 Bayesian Non-Parametric Methods

While the methods above require the specification of a number of topics $K$, there exists a class of Bayesian nonparametric methods for topic modeling that do not require choosing the number of topics $K$ and instead infer $K$ from the data. These methods are based on the hierarchical Dirichlet process, which allows for the estimation of $K$ along with the other parameters[31]. Several extensions of this method have been developed[32], including methods to improve the computational efficiency of the algorithm using online variational inference[33]. The hierarchical Dirichlet process produces a prior over $K$[34], and thus the SEEK-VEC framework could be used to compare the range of plausible values of $K$. However, due to the computational expense of these methods and the lack of off-the-shelf software packages that limit their practical utility, we do not consider them in full here.

##### A.2.3 Extended Simulation Results

Supplemental Fig. A3 shows extended simulation results for diagonal correlation and off-diagonal correlation, respectively. Under the strong anchor condition, rows represent different values of the threshold $t$. Under the weak anchor condition, rows represent different anchor word scale factors ($sf$), with smaller scale factors representing weaker anchor words. The loading of an anchor word is $sf$ times the loading of a non-anchor word. For a pure anchor word, non-anchor words have a loading of 0 and anchor words have a loading of 1; therefore, the higher $sf$ is, the closer the anchor words resemble pure anchor words. These figures additionally include the results for SEEK-VEC with a variety of upper limits for the number of topics included. For example, The model $O7$ ensembles $K = 3$ to $K = 7$.

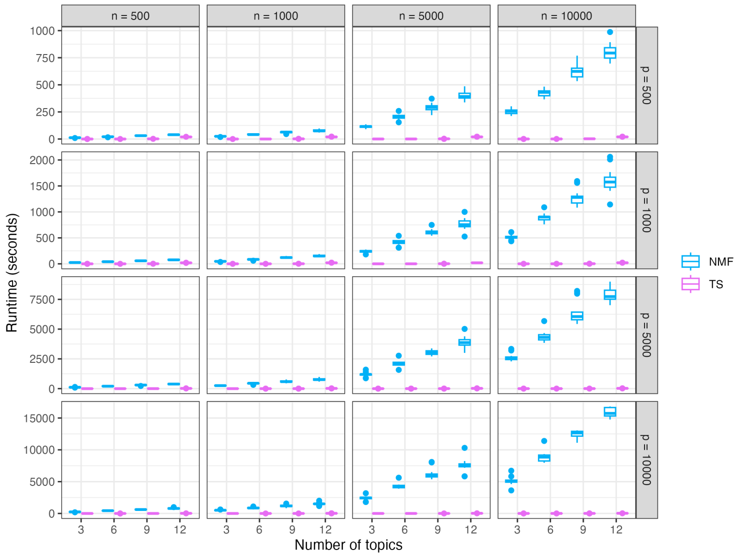


Supplemental Figure A2: Simulation results comparing the runtime of fastTopics (“NMF"), and TopicScore (“TS"). Columns represent varying numbers of documents and rows represent vocabulary size.

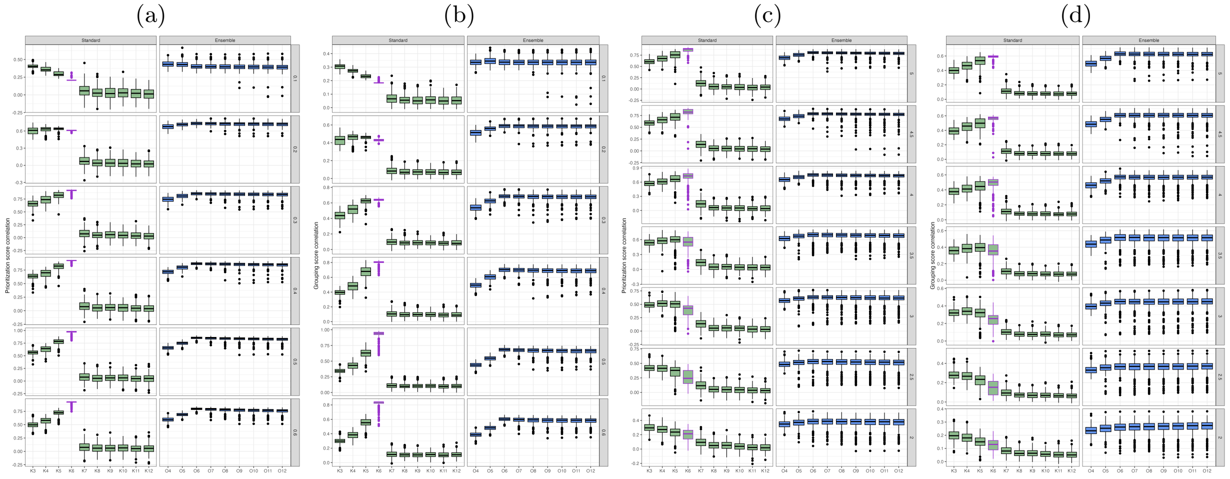


Supplemental Figure A3: Model $KX$ represents a standard topic model with $K = X$ topics. Model $OX$ represents the ensemble method using $K = 3, ..., X$ topics. (a-b) Prioritization score and grouping score correlations under the strong anchor condition across varying threshold levels. (c-d) Prioritization score and grouping score correlations with $t = 0.3$ and varying anchor word scale factors: as the anchor scale word factor decreases, the loadings of anchor words become more similar to the loadings of non-anchor words. Specifically, an anchor word scale factor of 5 represents a setting where the loading of an identifying or metainformative word is 5 times that of an uninformative word.

##### A.2.4 Comparison with alternate averaging strategies

Supplemental Fig. A4 shows the equivalent of Fig. 2c for the alternate ensemble strategies of uniform averaging (Supplemental Fig. A4a) and averaging of matrices normalized by their Frobenius norm (Supplemental Fig. A4b). Compared to SEEK-VEC, which implements eigenscore-weighted averaging, these methods are much more sensitive to hyperparameter selection.

##### A.2.5 Effect of $K$ Misspecification on Topic-SCORE Simplex Embedding

Supplemental Fig. A5 shows the impact of the potential misspecification of $K$ with respect to the simplex structure and vertex hunting steps in Topic-SCORE. The data are simulated from $p = 100$ vocabulary words on $K = \{2, 3, 4\}$ topics, and for each, Topic-SCORE is run under the assumption that there are 3 topics. The rows of each vocabulary-topic matrix $\mathbf{A}$ are simulated to comprise three types of words: identifying words, metainformative words, and uninformative words. Identifying words have a relatively high weight on only one topic and a low weight on all other topics; they are pseudo-anchor words. Metainformative words have a relatively high weight on two topics and a low weight on the others; these words provide information on how topics are linked, as will be discussed in further sections. Uninformative words have approximately equal weights on all topics, and therefore do not contribute to the understanding or identification of any topics.

The center panel of Supplemental Fig. A5 shows the simplex embedding of a correctly specified topic model, where the true number of topics matches the inputted value of $K$. The pink triangles represent the three vertices found by the vertex hunting step, and the identifying words lie at these vertices as desired. The metainformative words lie along the edges of their respective topics, and the uninformative words lie in the center of the simplex. The left panel shows the embedding of an over-specified topic model. The embedding shows uninformative words in between the sets of identifying words (note that there cannot be metainformative words in a two-topic model), but there is no obvious simplex structure and vertex hunting step has incorrectly placed the vertices along the vertical spread of the uninformative words. Finally, the right panel shows an under-specified model, where the true number of topics is larger than the $K$ provided to Topic-SCORE. Again, the simplex structure is unclear, and two of the three vertices are not close to identifying words.

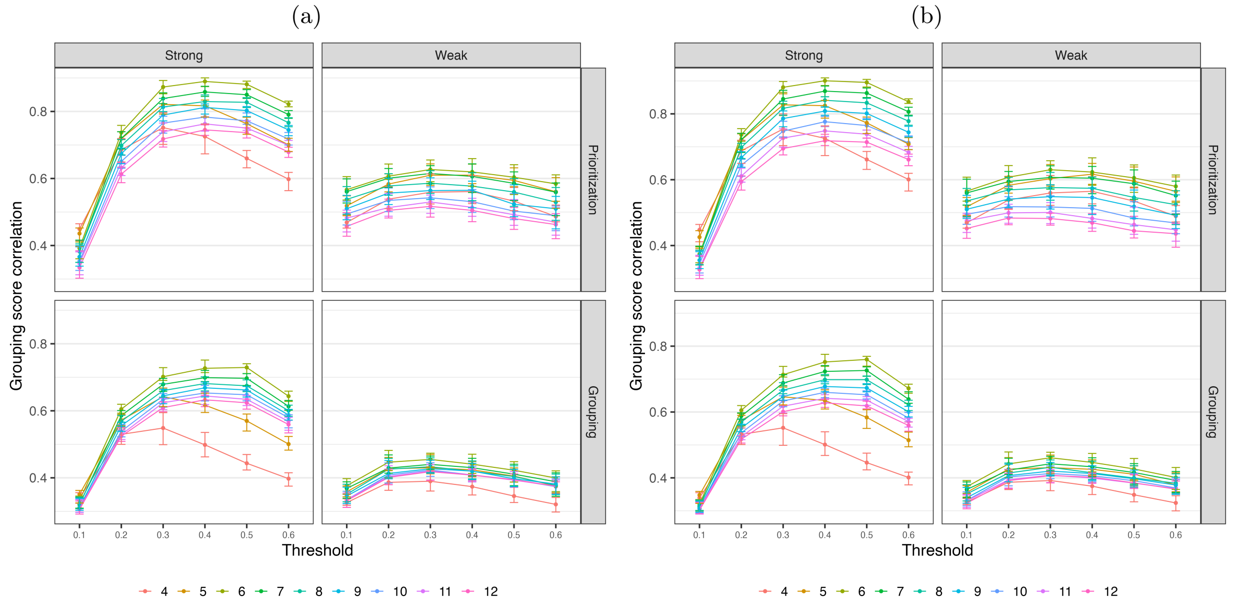


Supplemental Figure A4: Results across choice of upper limit for ensembling using the strategies of uniform averaging (left) and Frobenius normalized averaging (right).

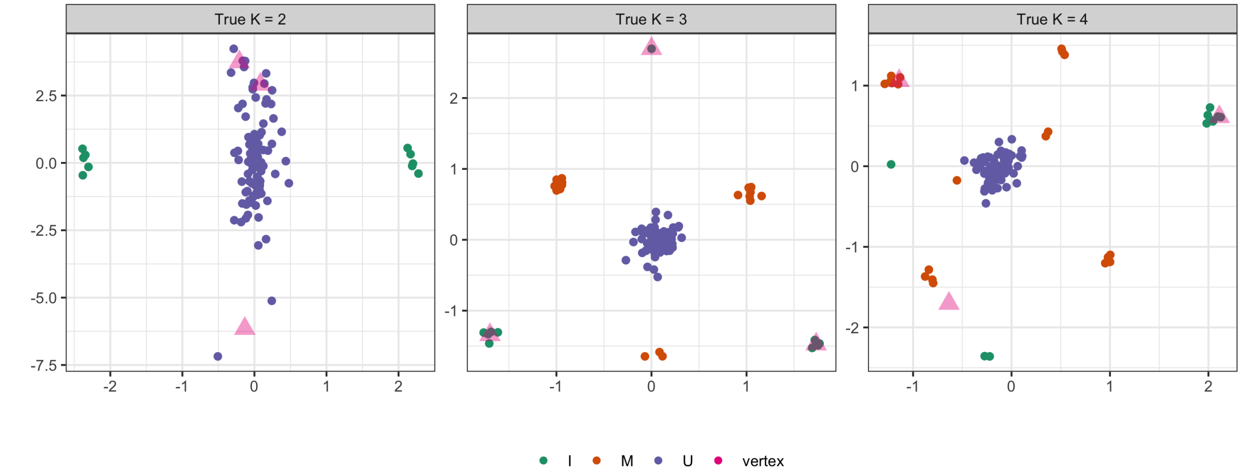


Supplemental Figure A5: Simplex embeddings and vertices for three models under the assumption that $K = 3$. The figure on the left shows the embedding of a model where the true number of topics is 2 (the topic model is overspecified). The figure in the center shows the embedding of a correctly-specified model. The figure on the left shows the embedding of a model where the true number of topics is 4 (the topic model is underspecified). The vocabulary words are colored by type: **I**dentifying (associated with only one topic), **M**etainformative (associated with two topics), and **U**ninformative (equally associated with all topics). Note that metainformative words cannot exist in the two-topic model.

###### A.2.6 Prioritization scores and grouping scores assess DSM-5 subscales in the SPARK Adult Self-Report

The SPARK study from the Simons Foundation Autism Research Initiative is an ongoing research study with over 100,000 participants with autism.[35] One source of phenotypic data is the Adult Self-Report (ASR), a well-validated survey instrument that is part of the Achenbach System of Empirically Based Assessment and measures adult psychopathology.[36] For the ASR, individuals are asked to rate themselves on 123 items, with the scoring options of *0 = Not True*, *1 = Somewhat or Sometimes True*, and *2 = Very True or Often True*. Of these items, 69 of the items on the ASR correspond to one of six DSM-5-Oriented Scales: Depressive Problems, Anxiety Problems, Somatic Problems, Avoidant Personality Problems, Attention Deficit/Hyperactivity Problems, and Antisocial Personality Problems. Items on these scales were rated to have strong correspondence to DSM-5 diagnostic criteria for particular disorders or categories of disorders.[37] Taking the items as the vocabulary and the individuals as documents, we can analyze the responses on these items from the 1,809 respondents with a confirmed autism diagnosis and assess how well these scales align with behavioral patterns among adults with autism.

Supplemental Fig. A6a shows the heatmap corresponding to $\widehat{\mathbf{\Phi}}_{t=0.5}$. We choose a high threshold value in this context because each item is designed to contain signal relating to an underlying pattern, namely the corresponding DSM-5 scale. The heatmap is filtered to rows containing a grouping score of 0.5 or greater, representing a weighted majority consensus of a significant relationship among items. Four blocks emerge: a Somatic block, a small Avoidant block, and two mixed and overlapping ADHD/Antisocial blocks. The mixed blocks align with research showing that individuals with ADHD are more likely to engage in antisocial behavior.[38,39] No items on the Depression or Anxiety subscales have a grouping score of 0.5 or greater with any other item, indicating that they do not contain strong signal of underlying response patterns in a population of individuals with autism. Notably, there is a lack of persistent shared signal despite choosing a very high threshold of $t = 0.5$; the Depression and Anxiety subscale items are not being co-identified as hallmark terms despite a very permissive threshold. While individuals with autism are four times more likely to experience depression than individuals without autism,[40] current diagnostic criteria for depression may not generalize to individuals with autism, for whom depression may present differently.[41] Similarly, while an estimated 50% of individuals with autism experience anxiety, anxiety may have an atypical presentation in

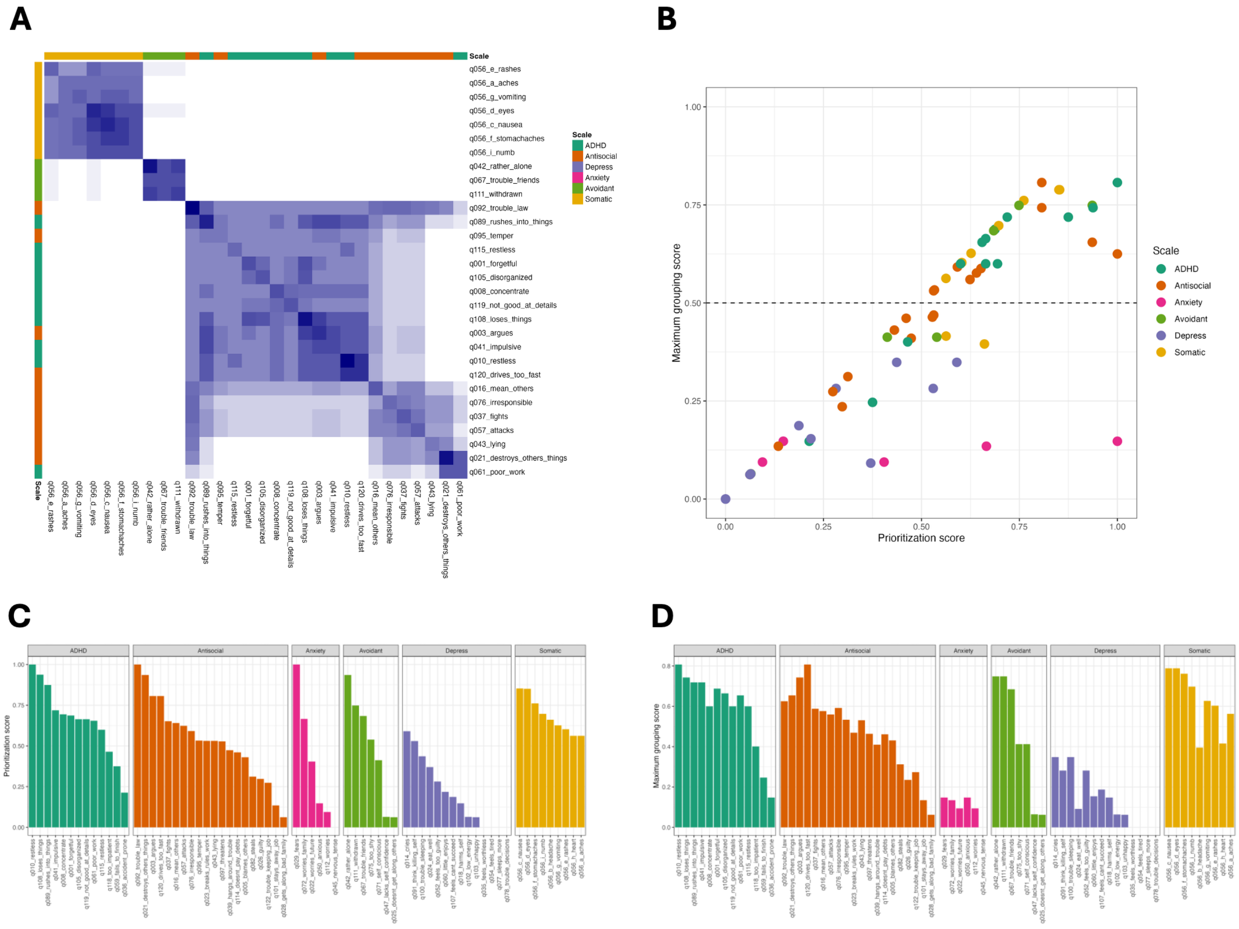


Supplemental Figure A6: (a) Heatmap of $\widehat{\mathbf{\Phi}}_{t=0.5}$. Items are annotated by their corresponding DSM-V subscale. (b) Scatterplot showing the patterns of prioritization versus maximum grouping scores in $\widehat{\mathbf{\Phi}}_{t=0.5}$. The dashed lined indicates the filtering threshold for plotting in (a). As seen in (a), all Depression and Anxiety items fall below the threshold. (c) Plots of the prioritization scores of each item in $\widehat{\mathbf{\Phi}}_{t=0.5}$. (d) Plots of the maximum grouping scores of each item in $\widehat{\mathbf{\Phi}}_{t=0.5}$.

these individuals that makes standard diagnostic criteria less effective.[42]

Supplemental Fig. A6b shows a scatterplot comparing each item's prioritization score and maximum grouping score, which allows for a classification of the survey items. First, many items fall along the diagonal. These survey items have at least one other item with which they are consistently co-identified, regardless of signal strength. For items falling below the diagonal, this means that their co-identification is less consistent: there is no other item with which they are consistently paired. As an example, we can see that *q_092_trouble_law* ("I do things that may cause me trouble with the law") has a prioritization score of 1 (Supplemental Fig. A6c), indicating that it is consistently identified as a hallmark item to an underlying latent pattern of responses. However, its maximum grouping score is less than 1 (Supplemental Fig. A6d), meaning it is indicative of different patterns in different models. This can be seen visually in Supplemental Fig. A6a, where it shows moderately strong overlap with both the majority-ADHD and majority-Antisocial blocks. Another interesting case is *q_029_fears* ("I am afraid of certain animals, situations, or places"), which has a prioritization score of 1 but a very low maximum grouping score. This shows that this item is consistently identified as the only hallmark item in a topic. Here, SEEK-VEC provides useful exploratory insight for survey design: this question seems to be ascertaining a concept that is distinct from the other items, and therefore may be misleading if used in a composite score. Examining the prioritization and grouping scores of these items provides key insight into response patterns and appropriate instrument design for assessing DSM-5 categories in adults with autism.

##### A.2.7 Meta-structure matrix reveals stochastic instability in BERTopic

BERTopic is a popular transformer-based topic modeling method that leverages machine learning and natural language processing principles to derive topic clusters from document embeddings.[24] BERTopic leverages UMAP,[43] a nonlinear dimension reduction method, prior to clustering the document embeddings. As UMAP is a stochastic method, we used SEEK-VEC to explore the stability of the BERTopic output with respect to this stochasticity.

We ran BERTopic five times on the *20 newsgroups* dataset, which contains approximately 18,000 text documents on 20 topics.[28] For each run, we used the default BERTopic parameters, specifying only the random seed (via *random_state*) in the underlying UMAP dimension reduction. Supplemental Fig. A7 shows the stability of the top 10 words from the top 10 topics from one of the models with respect to the set of five models. Several interesting patterns emerge. First, Topic 1 appears to be fully stable, with the top words consistent across random states. Topics 2, 3, and 7 are also highly stable. However, many of the top words in Topics 4 and 5 appear to be co-identified as top words in other models, linking the topics. Further examination of these topics shows clear overlap in their content; Topic 4 words include "cars," "v8", and "convertible", Topic 5 words include "motorcycle" and "brake", and overlapping words include "miles", "engine", and "car." Further, while a few of the words in Topic 8 are consistently co-identified as top words for the same topic ("mouse", "windows", "mbytes"), the remainder of the words do not appear to be consistently identified as top words for the same topic across models. Overall, the SEEK-VEC stability analysis demonstrates that the results from BERTopic may be vulnerable to the underlying stochasticity of UMAP.

##### A.2.8 MADStat SEEK-VEC word blocks

Block 1: aberr, aoptim, array, balanc, block, cyclic, design, doptim, experiment, factori, fraction, latin, nonregular, optim, orthogon, prime, resolut, satur, twofactor, twolevel

Block 2: band, biascorrect, confid, coverag, credibl, edgeworth, interv, jackknif, narrow, percentil, pivot, plugin, shorter, stein, width

Block 3: algorithm, anneal, descent, expectationmaxim, fast, faster, gradient, kmean, learn, machin, metropoli, metropolishast, mine, nonconvex, onlin, realworld, scalabl, slow, speed, stateoftheart, supervis, svm, task, text

Block 4: acut, air, childhood, crossvalid, diabet, event, femal, highthroughput, histori, immun, infect, occurr, outbreak, overfit, retrospect, survivor, viral, visit

Block 5: backfit, coeffici, cook, curs, leastsquar, linear, logist, nconsist, ordinari, quantil, regress, regressor, rootn, singleindex, varyingcoeffici

Block 6: causespecif, censor, continuoustim, declin, failur, hit, intervalcensor, landmark, month, onset, period, renew, repair, semimarkov, seri, state, surviv, time, wait

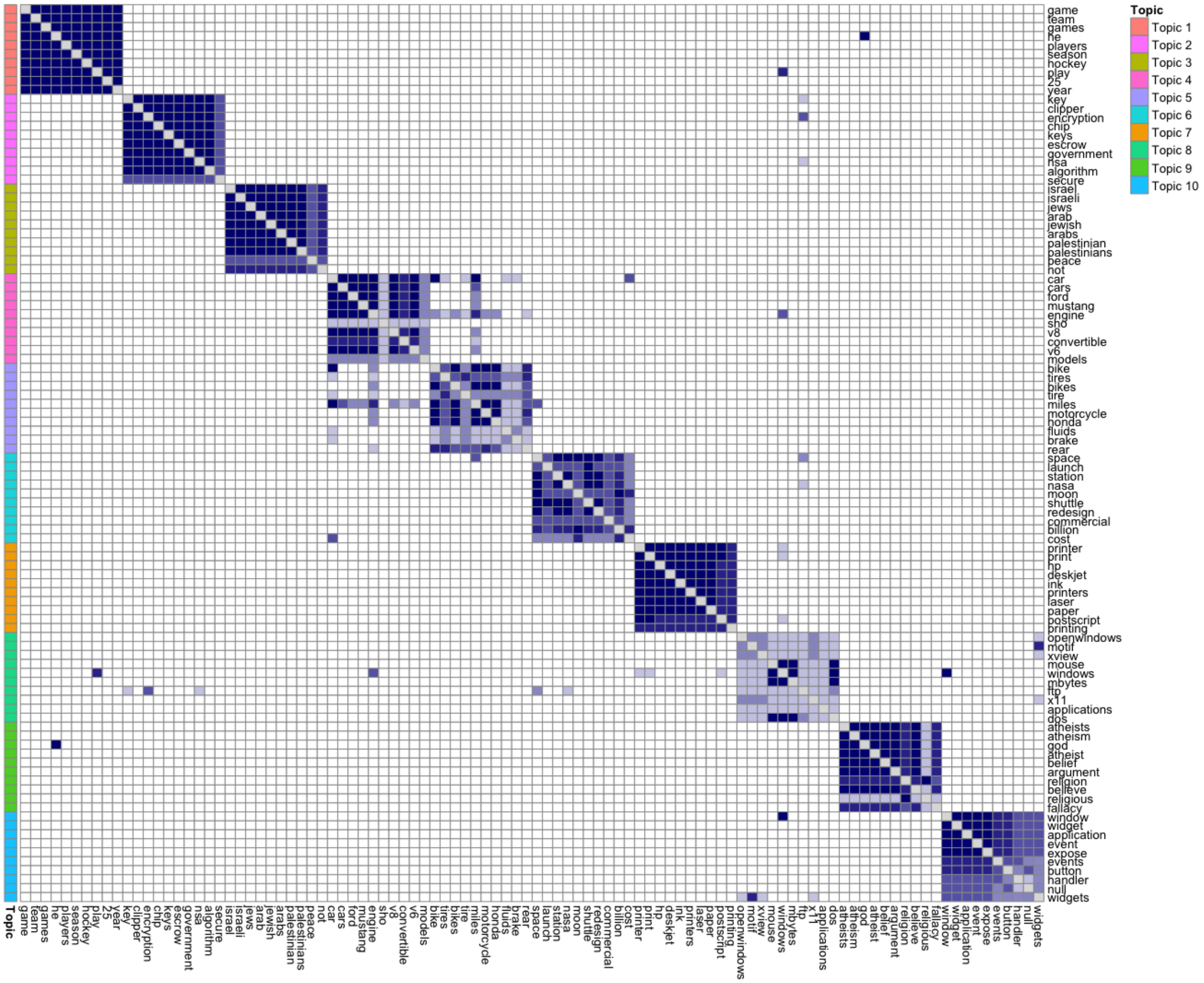


Supplemental Figure A7: Stability analysis of the top 10 topics derived from the 20 newsgroups dataset using BERTopic. Stability is assessed with respect to five identical BERTopic models, varying only the random seed used in the underlying UMAP procedure.

Block 7: antiretrovir, arm, assign, benefici, causal, clinic, clinician, complianc, depress, effect, imbal, intervent, metaanalys, metaanalysi, mixedeffect, noncompli, outcom, placebo, prognost, subgroup, symptom, therapi, timetoev, treatment, trial, vaccin, withinsubject

Block 8: corollari, ddimension, element, ergod, gumbel, inequ, infin, levi, nonneg, phi, probab, proof, theorem, tild, walk

Block 9: bay, bayesian, conjug, default, frequentist, improp, jeffrey, noninform, opinion, prio

Block 10: consecut, epsilon, inadmiss, lnorm, minimax, realvalu, regim, theta, typeii

Block 11: bonferroni, control, discoveri, fals, familywis, fdr, inflat, stepdown

Block 12: alzheim, diseas, genet, genomewid, phenotyp, polymorph

Block 13: concomit, explanatori, forest, instrument, manifest, variabl
